# Association Analysis of *LCORL* Genetic Variant (rs657074013A>AT) with Body-Height and Skeletal Frame in Pakistani Goats

**DOI:** 10.1101/2022.10.09.511453

**Authors:** Areeb Khalid, Hajra Ashraf, Hibba Asim, Maleeka Ayman, Rashid Saif

**Affiliations:** Decode Genomics, 323-D, Punjab University Employees Housing Scheme, Lahore, Pakistan

**Keywords:** Pakistani *Capra hircus*, 6:37928641, Large body-size, *LCORL* gene, Genetic marker

## Abstract

Goat is one of the most valued and cost-effective household animals serving humanity by providing its meat, milk, fiber, skin and manure. Particularly, meat and milk production depends on the physical traits of the animals e.g., large body-size/skeletal frame. Previous studies has shown that the *LCORL* gene is associated with the subject traits in humans and many other animal species. So, the current study is aimed to investigate the genetic variability/association of rs657074013A>AT variant within diverse goats from Pakistan. By using ARMS-PCR, a total of 51 samples were genotyped with allele-specific primer-set, results of the current study depicted that, 27% population is homozygous wild-type (A/A), 24% homozygous-mutant (AT/AT) and 49% is heterozygous (A/AT). Hardy Weinberg Equilibrium (*HWE*) analysis revealed that our overall sampled population obeys this principle with *χ*2(2, *N* = 51) = 0.046, *p* = 0.9730. Similarly, a significant genetic association was also envisioned by Chi-square statistics *p* = 9.60 × 10^−5^ using the PLINK data analysis toolset. Furthermore, alternative allele frequencies of 0.68 and 0.28 were also calculated within cases (n=25) and control (n=26) cohorts respectively along with an odds ratio of 5.242. As a nutshell, this pilot study improved our genomic insight by observing the variability and association of the subject *LCORL* genetic variant (c.828_829insA) in the Pakistani goat population, which may be used to improve this economically important trait by adopting marker-assisted breeding strategies and further investigated within other livestock species as well.

## Introduction

*Capra hircus*, commonly known as domestic goat, is a member of the animal family B*ovidae*. Approximately, 40 animal species are domesticated all over the world [1], and goats are one of the oldest species being domesticated as a vital producer of meat, milk, hair, fiber and skin. There are 37 goat breeds in Pakistan. Annually, goats produce around 275 thousand tonnes of meat, 851 thousand tonnes of milk, 25 million skins and 21.4 thousand tonnes of hair and thus play an important role in the economy nationwide [2]. Goat meat is one of the most desired around the world. Due to rising demand for goat meat, farmers are interested in raising and breeding goats for more protein yield. So, body-height and skeletal-frame is one of the targeted trait to improve meat production through adopting marker-assisted breeding strategies.

The main factor that control animals’ height and skeletal stature is their genetic makeup as well as the environmental factors. There are plethora of genes that control body-height and skeletal-frame in different animal species, such as the Sperm Associated Antigen-17 (*SPAG17*) and Pleomorphic Adenocarcinoma (*PLAG1*) genes [3, 4]. But the gene that has been the most studies and observed to be strongly associated with frame-size of goats is the Ligand Dependent Receptor Co-repressor Like (*LCORL)*, also known as the Mblk-1-related protein [5]. *LCORL* encodes a transcriptional factor that has been linked to body-stature and skeletal-frame. In large-sized goats, the truncation may interfere with *LCORL*’s transcription factor binding to its target, because the N-or C-terminal region of this protein is removed by proteolysis, premature protein elongation occurs due to the gain of stop codon and a nonsense mutation in the structured gene, it also has a function in the process of spermatogenesis.

In *Capra hircus*, the *LCORL* gene has been mapped on Chr.6 (NC_030813.1) genome assembly ARS1.2 (GCF_001704415.2) having 8 exons in total [6]. It has been identified in previous studies that the c.828_829insA locus at genomic position 6:37928641, (rs657074013A>AT) with r.1225 nucleotide position of transcript ID: (XM_018049322.1) located on exon 7 (Figure 4), which is found highly variable with goat-height and skeletal-frame [5]. The aim of the current study is to evaluate the association of the aforementioned variant with heighted phenotypic trait in Pakistani goat population for improving the overall meat and milk production in this valued species, at the same time, the subject trait may be improved in other animal species as well by investigating the same variant in other livestocks species.

## Materials and Methods

### Sample collection and DNA extraction

Blood samples of 51 goats were collected to evaluate the genetic association between the *LCORL* variant (rs657074013A>AT) with their body-height and skeletal-frame. For sampling, two goat groups were made, one is categorized as a heighted-cohort (n = 25, wither-height ≥ 36”) and the other one as a control-cohort (n = 26, wither-height < 36”) of the age bracket 1.5-2 years (Figure 1). EDTA vacutainers were used to collect and store blood samples at −20°C till further usage. GDSBio (https://www.gdsbio.com/en/) genomic DNA extraction kit was used to extract DNA from goat blood samples by following the manufacturer’s instructions.

**Figure 1:**
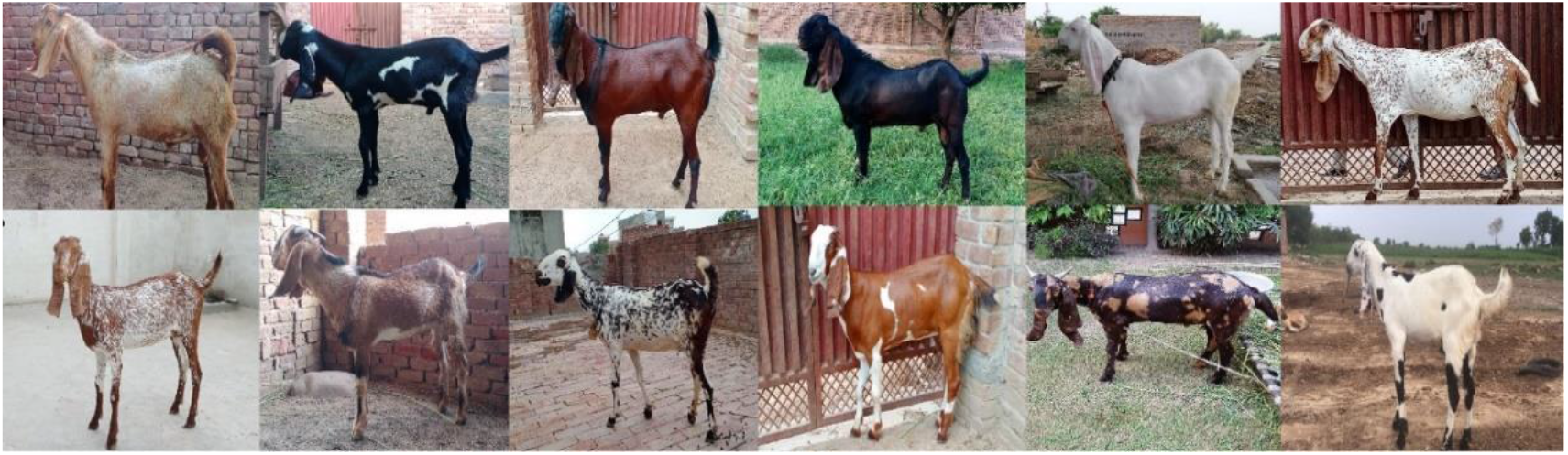
Few of the sampled goats, upper row animals are of wither-height ≥ 36”, while lower ones are below this threshold.

### Primer designing

ARMS-PCR primers were designed using OligoCalc software (http://biotools.nubic.northwestern.edu/OligoCalc.html) for the amplification of wild-type and mutant allele at c.828_829insA locus in goats against the transcript ID: XM_018049322.1. A total of five primers were designed, which were labelled as reverse common, forward normal and forward mutant ARMS primers with amplicon-size of 300 bp. Similarly, two internal control (IC) primers were also designed to amplify a region for PCR fidelity (Table 1).

**Table 1:**
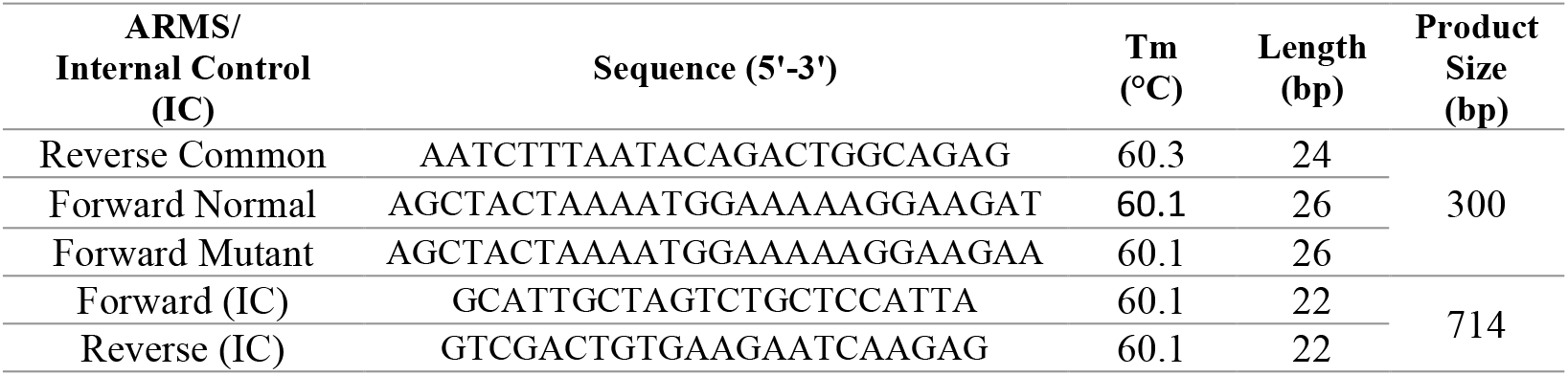
Primers sequences with its attributes

### DNA amplification

The ARMS-PCR reaction was carried out using a SimpliAmp thermal cycler (Applied Biosystems). Two PCR reactions were carried-out separately with each sample having normal (N) and mutant (M) type ARMS allele specific forward primer along with reverse common primer. Simultaneously, two regular primers were also used to amplify the genomic region as an internal control in few of the random samples. A total of 16µL of the reaction mixture was prepared consisting 2µL of 50ng/µL genomic DNA, 10mM of each primer, 0.05IU/µL of *Taq* polymerase, 2.5mM MgCl2, 2.5mM dNTPs, 1x *Taq* buffer and PCR-grade water. The PCR protocol was adopted with 5-minutes of initial denaturation at 95°C followed by 30 cycles of denaturation (95°C for 45 sec.), annealing (60°C for 30 sec.), extension (72°C for 45 sec.) with the final extension at 72°C for 10-minutes and stored at 4°C (Figure 2).

**Figure 2:**
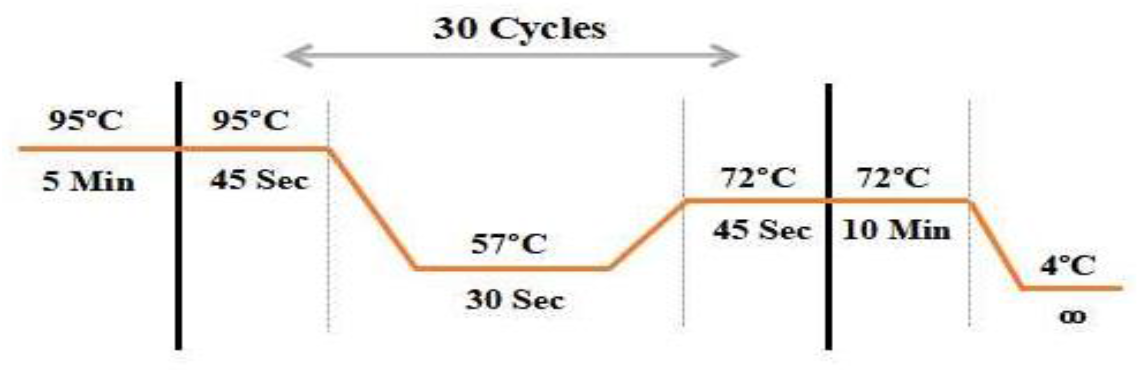
Thermal cyclic conditions of ARMS-PCR

### Statistical analysis

Hardy Weinberg Equilibrium (*HWE*) *was* used to calculate the observed & expected allelic and genotypic frequencies by obeying *p*2 + 2*pq* + *q*2 = 1 equation and further Chi-square analysis was also conducted using 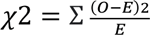 equation to calculate the *p* = value to check whether our sampled population is in accoradance with *HWE* or not.

Secondly, PLINK data analysis toolset was used to evaluate the association of subject variant with the body-height phenotype after screening the rs657074013A>AT locus in the sampled population of 51 goats. Alternative allele frequencies in cases and controls along with *p*-value and odds-ratio were also calculated in overall goat population.

## Results

In the current study, *LCORL* variant (rs657074013A>AT) showed the variability in Pakistani goat population, which is in accordance to the other goat populations of the world, and depicts that, this Chr.6 locus 6:37928641 is under-selection in large-sized Pakistani goat breeds.

A total of 51 samples were genotyped, (cases=25, wither-height ≥ 36”) and (controls=26 wither-height < 36”). After experimental and statistical analysis, it was concluded that, there are 2 homozygous wild-type (A/A), 11 homozygous-mutant (AT/AT) and 12 heterozygous (A/AT) individuals were observed among the heighted cohort. Similarly, 12, 01 and 13 goats are homozygous wild-type, homozygous-mutant and hetrozygous respectively in the control group. Hence, the overall genotypic frequency of homozygous wild-type in our sampled population is 0.27 (27%), homozygous-mutant is 0.24 (24%) and heterozygous is 0.49 (49%). Subsequently, both alleles (A)/(AT) frequencies were calculated as 0.52 and 0.48 respectively in oveall population. Thereafter, Hardy-Weinberg Equilibrium (*HWE*) Chi-square analysis was conducted to verify, whether our sampled population is obeying this principle or not with the following outcomes of *χ*2(2, *N* = 51) = 0.0546, *p* = 0.9730 which manifests that our population is in accordance with the *HWE* equilibrium as the *p*-value is above the set threshold confidence interval of 0.05 so accepting our null-hypothesis of observing *HWE*.

Moreover, genetic association analysis was conducted using PLINK data analysis toolset which demonstrated that the alternative allele frequency are 0.68 & 0.28 within our cases & control cohorts having *χ*2 statistics value of 15.65 and *p* − *value* = 9.60 × 10^−5^ showing a significant association of the screened variant with body-height phenotype in Pakistani goats. Likewise, 5.242 odds-ratio (OR) was also observed showing the prevelance of odds/mutants is almost 5-times heigher in cases vs controls. (Table 2).

**Table 2:**
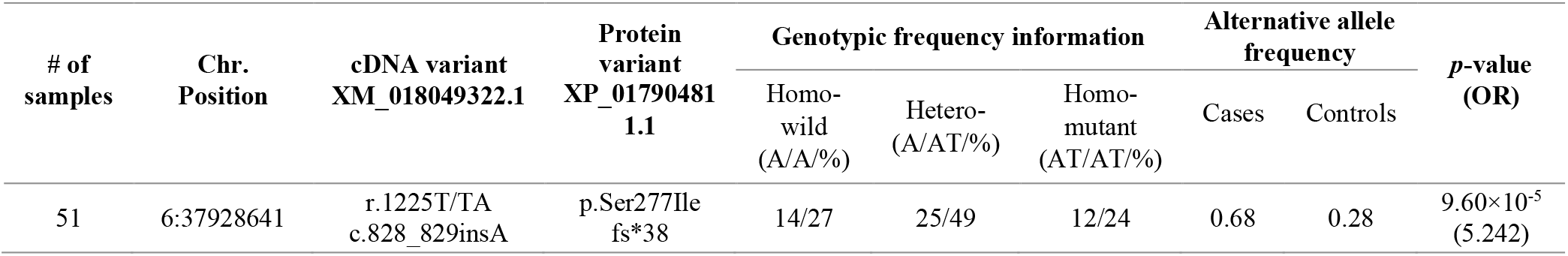
Plink association of subject variant with goat-wither height from Pakistan

This single base insertion variant of c.828-829insA causes frame-shift in LCORL protein and alters Serine amino acid (aa) to Isoleucine at 277^th^ position in this polypeptide which terminated this protein at 38^th^ position after 277^th^aa and resulted in 314 aa truncated LCORL potein instead of its wild-type protein of 1864 aa. This frame-shift mutation truncated almost ∼83% of the wild-type open reading frame due to gain of a premature stop codon at (ser277Ile fs*38) [5, 7]. This truncation may be responsible for the goat’s height and appeared as gain/loss of function mutation. Further, fuctional genetics studies are still needed to confirm and validate this postulated hypothsis.

## Discussion

The central importance and rising demand of meat is undeniable in the present world. Meat consumption has taken on a new vehemence especially in islamic countries such as Pakistan, where especially the goat meat is highly preffered and part of the daily diet of the masses. Moreover, goat milk is also religious preffered among the muslim communities to follow the Holy Prophet “Sunnah” with health benefits. So, goat rearing in Pakistan is highly profitable business and farmer are greatly interested in the goat farming and better performing herds [2]. So, the body-height and stature trait chosen in this study to meet he rising demand of goats in Pakistan. According to one of the genome-wide genomic selection signature study, the *LCORL* gene variant rs657074013A>AT on chr.6 was identified as the most relevant quantitative trait locus for height in goats [5].

The molecular mechanism underlying how the *LCORL* locus modulates body-size and height across several mammalian species is yet not clear [5]. In the current investigation, we tried to explore how the genetic variation influenced the subject trait in Pakistani goat poplation. Our findings revealed that there are only 2 & 12-homozygous wild-type, 11 & 01 homozygous-mutant and 12 & 13 goats are heterozygous in the cases and controls respectively, which depeicst the moderate fixation of mutant allele/genotype in the cases while heterozygous individuals are almost same in both groups that shows the random mating and locus hitchhiking in the population. Moreover, multiple sequence alignment of the subject locus is performed in nine mammalian species to check the conserved status of the locus which showing (T) nucleotide in goat as compare to (G) in horse, pig, humans and cattle while (C) in mouse (Figure 4).

**Figure 3:**
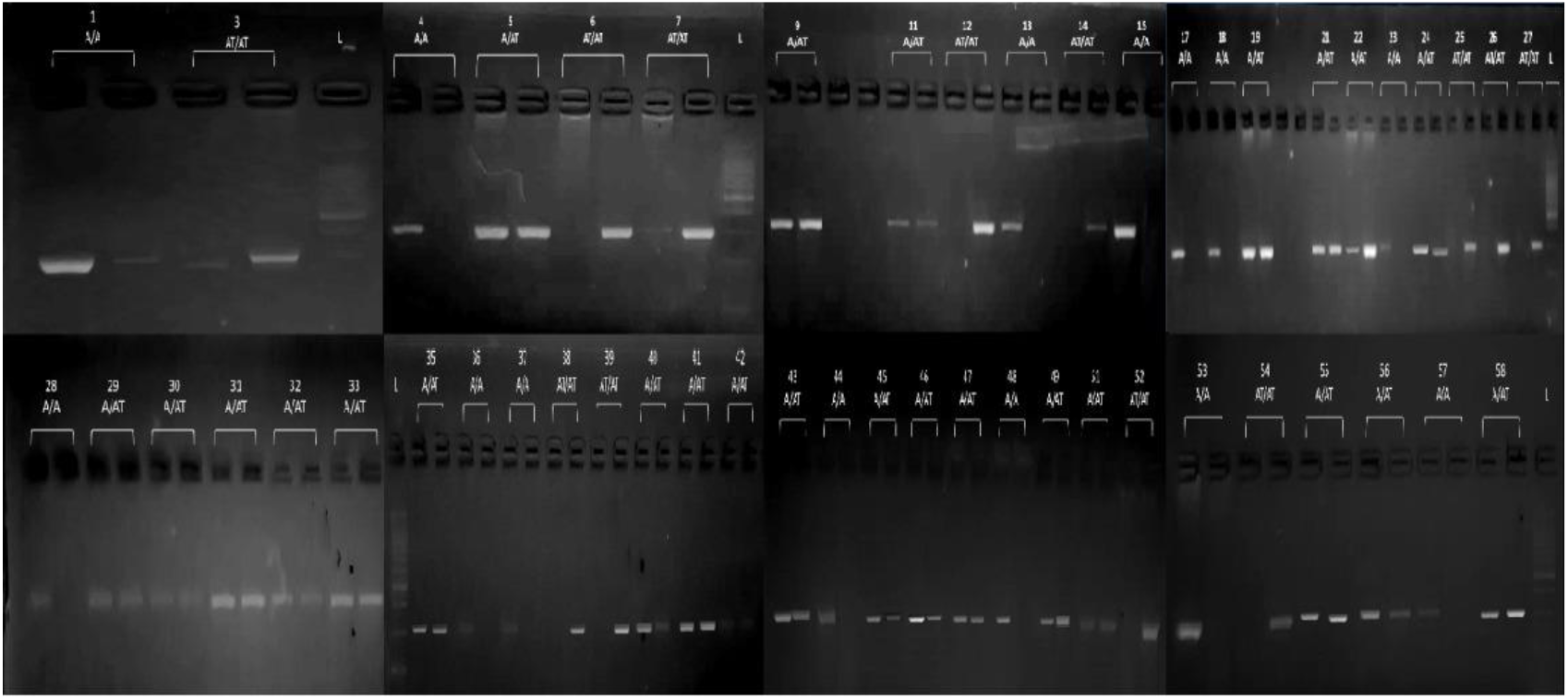
ARMS-PCR amplification of targeted variant within 51 sampled goats

**Figure 4:**
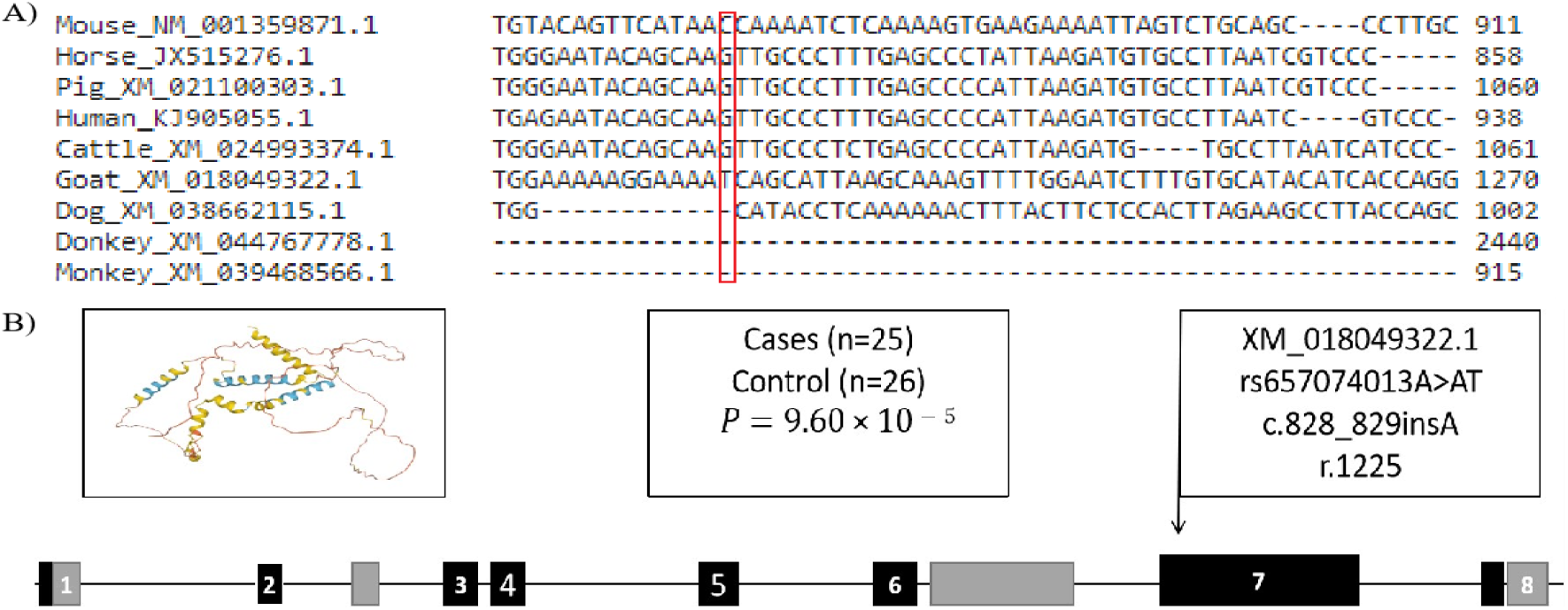
**A)** Multiple sequence slignment of genotyped variant in different species **B)** Genomic location of the subject variant and *LCORL* protein structure are shown

Previous studies have enlightened that *LCORL* is not only associated with body-stature in goats but in several other species as well e.g., cattle, sheep, horses, dogs, pigs, donkey and humans are also associated with the same gene and its surrounding gene of *PLAG1* [4, 8-14]. In 2017, a study from China revealed the association of *LCORL* candidate gene with the bovine development and carcass traits [6]. Two other studies in 2019-20 found *LCORL* contributes in body-height and stature in sheep as well [9, 15]. Another German study from 2013 showed that the *LCORL/NCAPG* locus have significant contribution in the heighted trait in German warm blooded equines [16].

Furthermore, **t**he whole genome sequencing of *Canids* revealed many genomic regions including *LCORL* that affect the dog’s skeletal-frame and other morphological traits [11]. A study on domestic pigs in 2012 showed many selection signatures harbour *LCORL* and its surrounding genes [12]. Similarly, a recent analysis in China found that the donkey’s body-size has also been attributed to the 3:112664848 locus in *LCORL* [8]. So, the current study also provide the enough evidence that the same gene is associated with the large-sized Pakistani goats which is in conformity to the other animal species of the world.

## Conclusion

The *LCORL* gene variant (rs657074013A/AT) is found to be remarkably associated with the wither-height in Pakistani goat population with *p* = 9.60 × 10^−5^ using PLINK association test. This variant may be validated further with genetic functional studied and can be used in marker assisted breeding strategies to improve the height and stature of the goat herds in Pakistan.

## Ethical Statement

For this research, ethical guidelines and permission are omitted. There was no special need for approval from an animal ethics committee. Sampled goats belong to the local farmers those who were approached through personal contacts and blood samples were taken upon their consent by obeying the animal handling ethics.

## Acknowledgement

Authors are obliged to the goat farmers for their support to provide blood samples.

## Authors Contribution

Areeb Khalid (AK), Hajra Ashraf (HA), Hibba Asim (HA) and Maleeka Ayman (MA) has equal contribution as first author, they all helped in blood samples collection, performed wet-lab and wrote the initial draft; Rashid Saif (RS) designed and envisaged the research project, trouble shooting in wet & dry-labs, arranged samples, applied statistical testing, editing, proof reading and correspondence of the manuscript.

